# Validation and Topic-driven Ranking for Biomedical Hypothesis Generation Systems

**DOI:** 10.1101/263897

**Authors:** Justin Sybrandt, Ilya Safro

## Abstract

Literature underpins research, providing the foundation for new ideas. But as the pace of science accelerates, many researchers struggle to stay current. To expedite their searches, some scientists leverage hypothesis generation (HG) systems, which can automatically inspect published papers to uncover novel implicit connections. With no foreseeable end to the driving pace of research, we expect these systems will become crucial for productive scientists, and later form the basis of intelligent automated discovery systems. Yet, many resort to expert analysis to validate such systems. This process is slow, hard to reproduce, and takes time away from other researchers. Therefore, we present a novel method to validate HG systems, which both scales to large validation sets and does not require expert input. We also introduce a number of new metrics to automatically identify plausible generated hypotheses. Through the study of published, highly cited, and noise predicates, we devise a validation challenge, which allows us to evaluate the performance of a HG system. Using an in-progress system, MOLIERE, as a case-study, we show the utility of our validation and ranking methods. So that others may reproduce our results, we provide our code, validation data, and results at bit.ly/2EtVshN.

## 1 Introduction

Literature underpins research and provides the groundwork upon which scientists construct new ideas. But while the foundation of published knowledge grows [24], so does the difficulty in surveying it. Currently the National Library of Medicine adds about 2,300 papers a day to MEDLINE [30], and this is likely to increase given that scientific output doubles every nine years [50]. Working in the modern deluge of information, researchers can miss valuable connections.

*Undiscovered public knowledge* is the information no one explicitly knows, but has already been implicitly published [46]. For instance, in 1986 no one knew that fish oil was a treatment for Raynaud’s Syndrome [45]. Instead, we knew separately that fish oil decreases a number of factors, and that those same factors worsen Raynaud’s Syndrome. Arrowsmith, the software which uncovered this finding, was the first *hypothesis generation system* (HG) [39].

The importance of these systems rises alongside the pace of scientific output; an abundance of literature implies an abundance of implicit connections. Yet, while many projects focus on new ways to find potential discoveries [51, 49, 27, 10, 46, 2, 42], few have a way to rigorously verify the quality of the connections posed by their system [7].

HG systems are hard to verify because they attempt to uncover novel information, a challenging task even for human researchers. To understand whether a potential connection is novel, these systems must model novelty itself. While there are verifiable models for novelty in specific contexts, each is trained to detect patterns similar to those present in a training set, which is conducive to traditional cross-validation [14, 23]. Some examples include using non-negative matrix factorization [25] to uncover protein-protein interactions [12], or to discover mutational cancer signatures [3]. In other domains, groups are using Bayesian surprise [20] to identify interesting topological features [33] or unexpected measurements for the purpose of uncertainty quantification [4]. In contrast to all of the above, HG systems strive to detect patterns *absent* from training data (but not necessarily outliers), and in doing so inspire new inquiry.

Without a viable alternative, the HG community primarily turns to expert analysis in order to validate the output of these systems [7, 51, 49, 27, 10, 46, 2]. Many methods also require an expert’s input to rank their potential connections [49, 27, 46]. But, getting expert input is time consuming, introduces bias, and cannot scale.

**Our contribution:** We propose a novel method to thoroughly validate HG systems without expert input. Any system that provides a ranking criteria along with its generated hypotheses can leverage this validation method. In addition, we present a number of new ranking strategies that apply to any HG system that produces topic models. We implement each ranking strategy within our in-progress system, MOLIERE [49], in order to compare their performance with respect to our validation method. Although our work focuses on biomedical research, the methods we discuss easily generalize to HG in any domain.

Our methodology leverages predicates, or subject-verb-object statements, found within the MEDLINE dataset of over 27 million scientific abstracts. For simplicity and reproducibility we use the Semantic Medical Database (SemMedDB) [22], which defines predicates using identifiers provided by the Universal Medical Language System (UMLS) [1]. We identify the first publication date of each unique predicate in order to find new connections published each year. Because these connections all come from titles and abstracts of MEDLINE papers, we note that before a predicate’s first publication, our system is unaware of any direct connection between its subject and object. Yet, a newly introduced predicate in a title or abstract represents a fruitful connection [8], the sort we would like a HG system to predict.

To begin our mass validation effort, that to the best of our knowledge has not been performed by other general-purpose HG systems, we select a “cut date,” which determines the set of publications provided to our system. We define the set of predicates that first occur after the cut date as the “published” set, and we subset it to identify predicates that first occurred in “highly-cited” papers. Next we generate a set of “noise” predicates that are randomly constructed and have never been published. We then run queries for predicates in all three sets using our HG system, which allows us to understand our ability to recover important connections. From there we rank our results and plot Receiver Operating Characteristic (ROC) curves [14] in order to estimate the overall performance of our system.

We find that by training a polynomial combination of multiple ranking metrics, MOLIERE is able to distinguish published and noise predicates with an ROC area of 0.834, and is able to distinguish highly cited predicates from noise with an ROC area of 0.874. This result is especially significant given that the main validation methods available, to both our system and other similar systems (see survey in [49]), were expert analysis and replicating the results of others [7].

## 2 Background

HG systems attempt to extract novel ideas from existing research. In this section, we begin by providing an overview on these systems and continue to explain how we leverage two complimentary techniques for understanding large text corpora: Topic Modeling [6] and Word Embedding [29]. The former identifies fuzzy clusters of terms, which allow us to glimpse the main domains of discourse across a document set. The latter embeds terms into a real-valued vector space, which allows us to convey semantic relationships using various distance metrics.

### 2.1 Extracting Information from Hypothesis Generation Systems

Swanson and Smalheiser created the first HG system Arrowsmith [39], and in doing so outlined the ABC model for discovery [48]. Although this approach has limitations [36], its conventions and intuitions remain in modern approaches [42].

In the ABC model, users run queries by specifying two keywords *a* and *c*. The goal of a HG system is to discover some entity *b* such that there are known relationships “*a* → *b*” and “*b* → *c*,” which allow us to imply the relationship between *a* and *c*. Because many connections may require more than one element *b* to describe, researchers apply other techniques, such as topic models in our case, to describe these connections.

We center this work around our in-progress system, MOLIERE [49]. Once a user queries *a* and *c*, our system identifies a relevant region within our multi-layered knowledge network, which consists of papers, terms, phrases, and various types of links. We extract abstracts from this region and create a sub-corpus upon which we generate a topic model. This topic model describes groups of related terms, which we treat as a hypothesis. Until now, like other systems, we left the analysis of MOLIERE’s output to experts, but through the methods described in Section 4 we perform this analysis automatically.

### 2.2 Word and Phrase Embedding

The method of finding dense vector representations of words is often referred to as “`word2vec`”. In reality, this umbrella term references two different algorithms, the Continuous BOW (CBOW) method and the Skip-Gram method [29] Both rely on shallow neural networks in order to learn vectors through word-usage patterns.

MOLIERE uses `FastText` [21], a similar tool under the `word2vec` umbrella, to find high quality embeddings of medical entities. By preprocessing our text with the automatic phrase mining technique `ToPMine` [9], we improve these embeddings while finding multi-word medical terms such as “lung cancer” or “benign tumor.” We see in Figure 1 that `FastText` clusters similar biological terms, an observation we leverage to validate our HG system and derive a number of metrics.

**Figure 1:**
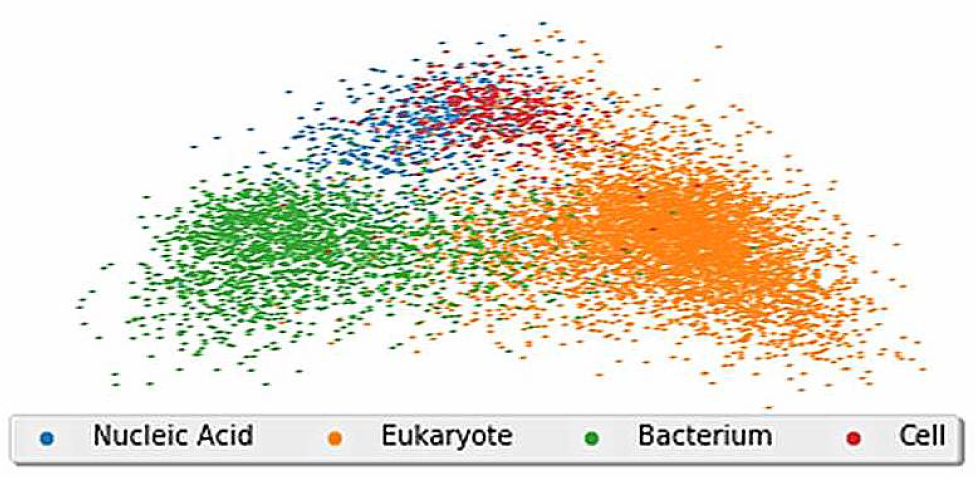
The above diagram shows a 2-D representation of the embeddings for over 8 thousand UMLS keywords within MOLIERE. We used singular value decomposition to reduce the dimensionality of these vectors from 500 to 2.

### 2.3 Topic Models

Latent Dirichlet Allocation (LDA) [6], the classical topic modeling method, groups keywords based on their document co-occurrence rates in order to describe the key “topics” of discourse. A topic is simply a probability distribution over a vocabulary, and each document from the input corpus is assumed to be a “mixture” of these topics. For instance, a topic model derived from New York Times articles would likely find one topic containing words such as “computer,” “website,” and “Internet,” while another topic may contains words such as “money,” “market,” and “stock.”

In the medical domain, we use topic models to understand trends across scientific literature. We look for groupings of entities such as genes, drugs, and diseases, which we then analyze to find novel connections. MOLIERE uses a parallel technique, PLDA+ [28] to quickly find topics from documents related to *a* and *c*. Because we pre-process MEDLINE articles with ToPMine, our resulting topic models include both words and phrases, which results in more meaningful topics.

### 2.4 Combing LDA and Word2Vec

There exists a symmetry between topic models and word embeddings [15]. The hidden layer of either CBOW or Skip-Gram captures semantic categories present in an input corpus. Each word in the vocabulary, when projected into the hidden layer, is represented as a mixture of hidden features — the same way we can view a word’s probability across a topic model.

A topic model is a weighted point cloud, where the embedding of each word is weighted by its probably within a given topic. Therefore, we can also represent the topic as a weighted centroid. In the following section we leverage both representations in order to mathematically describe the similarity between *a*, *c*, and each topic.

## 3 Validation Challenge

In order to unyoke automatic HG from expert analysis, we propose a challenge that any system can attempt, provided it can rank its proposed connections. A successful system ought to rank published connections higher than those we randomly created. So, we train a system given historical information, and create the “published,” “highly-cited,” and “noise” query sets. We pose these connections to an HG system, and rank its outputs to plot ROC curves, which determine whether published predicates are preferred to noise. Through the area under these ROC curves, a HG system demonstrates its quality at a large scale without expert analysis.

Our challenge starts with the Semantic Medical Database (SemMedDB) [22] that contains predicates extracted from MEDLINE defined on the set of UMLS terms [30]. For instance, predicate “C1619966 TREATS C0041296” represents a discovered fact “abatacept treats tuberculosis”. Because our system does not account for word order or verb, we look for distinct unordered word-pairs *a–c* instead. Using the metadata associated with each predicate, we note the date each unordered pair was first published. In Section 7 we discuss how we may improve our system to include this unused information.

From there, we select a “cut year.” For this challenge, we create our knowledge network [49] using only information published before the cut year. We then identify the set of SemMedDB unordered pairs *a–c* first published after the cut year provided *a* and *c* both occur in that year’s UMLS release. This “published set” of pairs represent new connections between existing entities, from the perspective of our HG system. We select 2010 as the cut year for our study in order to create a published set of over 1 million pairs.

Also, we create a set of “highly-cited” pairs by filtering the published set by citation count. We use data from SemMedDB, MEDLINE, and Semantic Scholar to create a set of 1,448 pairs from the published set that first occurred in a paper cited over 100 times. We intuit that this set is closer to the number of landmark discoveries since the cut-date, given that the published set is large and likely contains incidental or incorrect connections.

To provide negative examples, we generate a “noise set” of pairs by sampling the cut-year’s UMLS release, storing the pair only if it does not occur in SemMedDB. These pairs represent nonsensical connections between UMLS elements. Although it is possible that we may stumble across novel findings within the noise set, we assume this will occur infrequently enough to not effect our results.

We run *a–c* queries from each set through MOLIERE and create two ranked lists: published vs. noise (PvN) and highly-cited vs. noise (HCvN). After ranking each set, we generate ROC curves [14], which allow us to judge the quality of our HG system. If more published predicates occur earlier in the ranking than noise, the ROC area will be close to 1, otherwise it will be closer to 0.5.

## 4 Ranking Methods for Topic Model Driven Hypotheses

In order to rise to the challenge stated above, we advanced MOLIERE [49] to produce a numeric ranking that accompanies its resulting hypotheses. As described above, we use this ranking to calculate ROC curves and evaluate our system’s performance.

Another extremely important use case of ranking is related to massive query runs in hypothesis generation systems. Typically, biomedical researchers are interested in investigating connections of one-to-many types. For example, one disease can be queried versus all genes in order to establish what genes are related to it or one specific drug ingredient is verified versus all side effects. In practice, a researcher may be interested to run *a – c_i_* queries for tens of thousands *c_i_*’s. In such cases, ranking the hypotheses becomes vitally important.

In the following sections we present a number of promising approaches for ranking the topic model results from MOLIERE queries. The key intuition underpinning these metrics is depicted in Figure 2. That is, related objects are grouped together in vector space, so if *a* and *c* are related by a third entity *b*, we hypothesize *b*’s vector *ϵ*(*b*) ought to be similar to both *ϵ*(*a*) and *ϵ*(*c*). Because MOLIERE returns a topic model, rather than some single *b*, we quantify the potential relationship between *a* and *c* through their relationship to the topic model using the following metrics.

**Figure 2:**
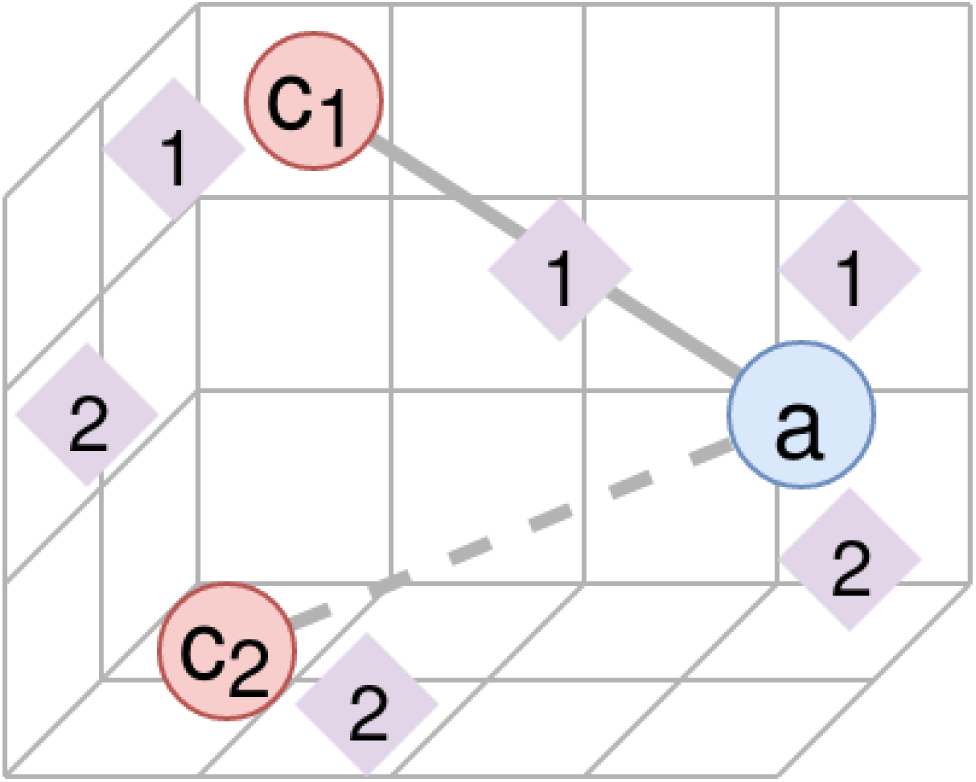
The above depicts two queries, *a–c*_1_ and *a–c*_2_, where *a–c*_1_ is a published connection and *a–c*_2_ is a noise connection. We see topics for each query represented as diamonds via Centr(*T_i_*). Although both queries lead to topics which are similar to *a*, *c*_1_, or *c*_2_, we find that the the presence of some topic which is similar to *both* objects of interest may indicate the published connection.

### 4.1 Similarity Between Query Words

As a baseline, we first consider two similarity metrics that do not include topic information: cosine similarity (CSIM) and Euclidean distance (*L*_2_), namely,

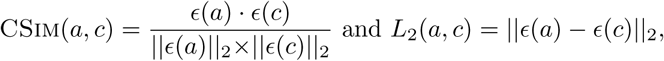

where *a* and *c* are the two objects of interest, and *ϵ*(*x*) is an embedding function (see Section 2.2). Note, when calculating ROC curves for the *L*_2_ metric, we will sort in reverse, meaning smaller distances ought to indicate published predicates.

These metrics indicate whether *a* and *c* share the same cluster with respect to the embedding space. Our observation is that this can be a good indication that *a* and *c* are of the same kind, or are conceptually related. This cluster intuition is shared by others studying similar embedding spaces [52].

### 4.2 Topic Model Correlation

The next metric attempts to uncover whether a and c are mutually similar to the generated topic model. This metric starts by creating vectors *v*(*a,T*) and *v*(*c,T*) which express each object’s similarity to topic model 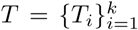 derived from an *a – c* query. We do so by calculating the weighted cosine similarity TOPICSIM(*x*, *T*_*i*_) between each topic *T_i_* and each object *x* ∈ {*a, c*}, namely,

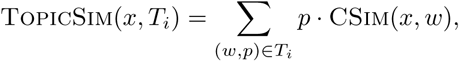

where a probability distribution over terms in *T_i_* is represented as word-probability pairs (*w, p*). This metric results in a value in the interval [-1, 1] to represent the weighted similarity of *x* with *T_i_*. The final similarity vectors *v*(*a,T*) and *v*(*c,T*) in ℝ^*k*^ are defined as

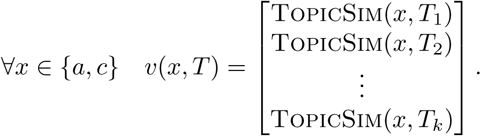

Finally, we can see how well *T* correlates with both *a* and *c* by taking yet another cosine similarity

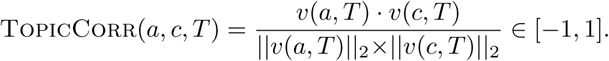

If TOPICCORR(*a, c, T*) is close to 1, then topics that are similar or dissimilar to *a* are also similar or dissimilar to *c*. This is supported by many experimental observations that if some explanation of the *a–c* connection exists within *T*, then we anticipate that many *T_i_* ∈ *T* would share these similarity relationships.

### 4.3 Similarity of Best Topic Centroid

While the above metric attempts to find a trend within the entire topic model *T*, this metric attempts to find just a single topic *T_i_* ∈ *T* that is likely to explain the *a – c* connection. This metric is most similar to that depicted in Figure 2. Each *T_i_* is represented in the embedding space by taking a weighted centroid over its word probability distribution. We then rate each topic by averaging its similarity with both queried words. The score for the overall hypothesis is simply the highest score among the topics.

We define the centroid of *T_i_* as

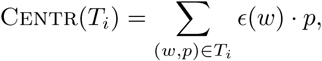

and then compare it to both *a* and *c* through cosine similarity and euclidean distance. When comparing with CSIM, we highly rank *T_i_*s with centroids located within the arc between *ϵ*(*a*) and *ϵ*(*c*). Because our embedding space identifies dimensions that help distinguish different types of objects, and because we trained a 500-dimensional embedding space, cosine similarity intuitively finds topics that share similar characteristics to both objects of interests. We define the best centroid similarity for CSIM as

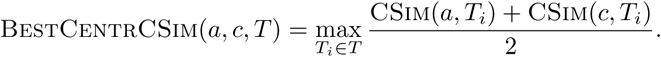

What we lose in the cosine similarity formulation is that clusters within our embedding space may be separate with respect to Euclidean distance but not cosine similarity. In order to evaluate the effect of this observation, we also formulate the best centroid metric with *L*_2_ distance. In this formulation we look for topics who occur as close to the midpoint between *ϵ*(*a*) and *ϵ*(*c*) as possible. We express this score as a ratio between that distance and the radius of the sphere with diameter from *ϵ*(*a*) to *ϵ*(*c*). In order to keep this metric in a similar range to the others, we limit its range to [0, 1], namely, for the midpoint *m* = (*ϵ*(*a*) + *ϵ*(*c*))/2,

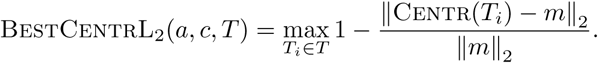

### 4.4 Cosine Similarly of Best Topic Per Word

In a similar effort to the above centroid-based metric, we attempt to find topics which are related to *a* and *c*, but this time on a per-word (or phrase) basis using TOPICSIM(*x*, *T_i_*) from Section 4.2. Now instead of looking across the entire topic model, we attempt to identify a single topic which is similar to both objects of interest. We do so by rating each topic by the lower of its two similarities, meaning the best topic overall will be similar to both query words.

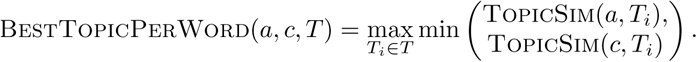

### 4.5 Network of Topic Centroids

A majority of the above metrics all rely on a single topic to describe the potential connection between *a* and *c*, but as Smalheizer points out in [37], a hypothesis may be best described as a “story” — a series of topics in our case. To model semantic connections between topics, we induce a nearest-neighbors network *N* from the set of vectors *V* = *ϵ*(*a*) ∪ *ϵ*(*b*) ∪ {CENTR(*T_i_*)}_*T*_*i*_ϵ*T*_ which form the set of nodes for *N*. In this case, we set the number of neighbors per node to the smallest value (that may be different for each query) such that there exists a path from *a* to *c*. Using this topic network, we attempt to model the semantic differences between published and noise predicates using network analytic metrics.

We depict two such networks in Figure 3, and observe that the connectivity between *a* and *c* from a published predicate is substantially stronger and more structured. In order to quantify this observed difference, we measure the average betweenness and eigenvector centrality [31] of nodes along a shortest path from *a* to *c* (denoted by *a* ~ *c*) within *N* to reflect possible information flow between *T_i_* ∈ *T*. This shortest path represents the series of links between key concepts present within our dataset that one might use to explain the relationship between *a* and *c*. We expect the connection linking *a* and *c* to be stronger if that path is more central to the topic network. Below we define metrics to quantify the differences in these topic networks. Such network analytic metrics are widely applied in semantic knowledge networks [41].

- TOPICWALKLENGTH(*a*, *c*, *T*): Length of shortest path *a* ~ *c*
- TOPICWALKBTWN(*a*, *c*, *T*): Avg. betweenness centrality of *a* ~ *c*
- TOPICWALKEIGEN(*a*, *c*, *T*): Avg. eigenvalue centrality of *a* ~ *c*
- TOPICNETCCOEF(*a*, *c*, *T*): Clustering coefficient of *N*
- TOPICNETMOD(*a*, *c*, *T*): Modularity of *N*.

**Figure 3:**
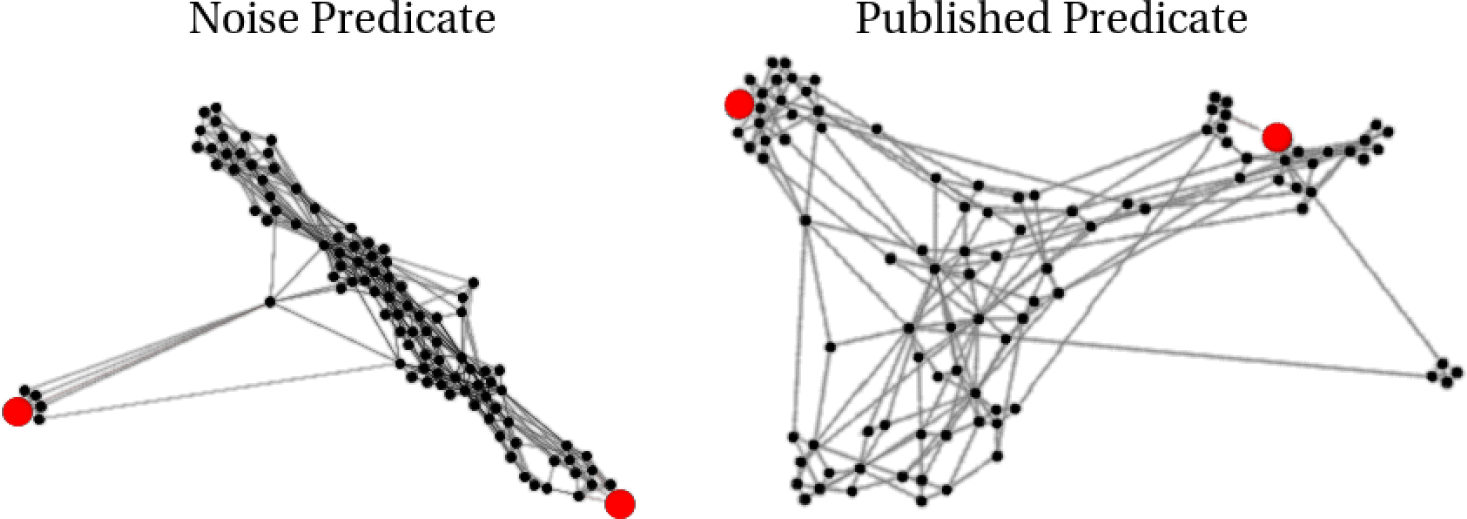
Above depicts two topic networks as described in Section 4.5. In this visualization, longer edges correspond to dissimilar neighbors. In red are objects *a* and *c*, which we queried to create these topics models. We observe that the connectivity between *a* and *c* from the published predicate is much higher than in the noisy example.

### 4.6 Combination of Multiple Metrics

Each of the above methods are based on different assumptions regarding topic model or embedding space properties exhibited by published connections. To leverage each metric’s strengths, we combined the top performing ones from each category into the following POLYMULTIPLE method. We explored polynomial combinations in the form of 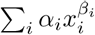 for ranges of *α_i_* ∈ [−1,1] and *β_i_* ∈ [1, 3] after scaling each *x_i_* to the [0,1] interval. Through a blackbox optimization technique, we searched over one-million parameter combinations, and present the best values Table 1. Omitted metrics were optimized with corresponding *α_i_* ≠ 0.

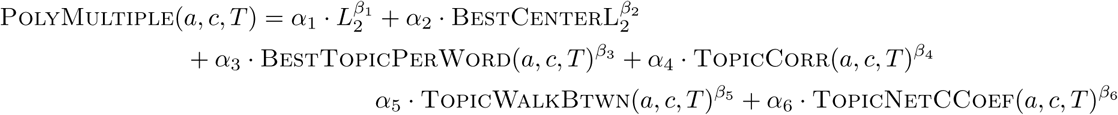

**Table 1:**
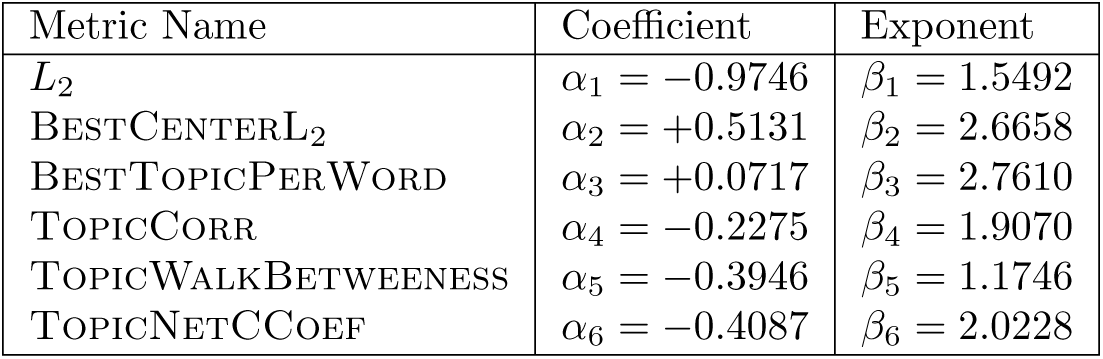
Best hyperparameters obtained via blackbox optimization for the PolyMultiple method. Note, each parameter was scaled to the [0, 1] interval before its input into the overall polynomial.

## 5 Results and Lessons Learned

As described in Section 3, our goal is to distinguish publishable connections from noise. We run MOLIERE to generate topic models related to published, noise, and highly cited pairs. Using this information, we plot ROC curves in Figures 4 and 5, and summarize the results in Table 2. These plots represent an analysis of 8,638 published vs. noise (PvN) pairs and 2,896 highly-cited vs. noise (HCvN) pairs (half of each set are noise). Although external factors limited the scale of our validation in both cases, we found that the resulting ROC areas are consistent for data sets of at least a few thousand pairs and different samplings.

**Figure 4:**
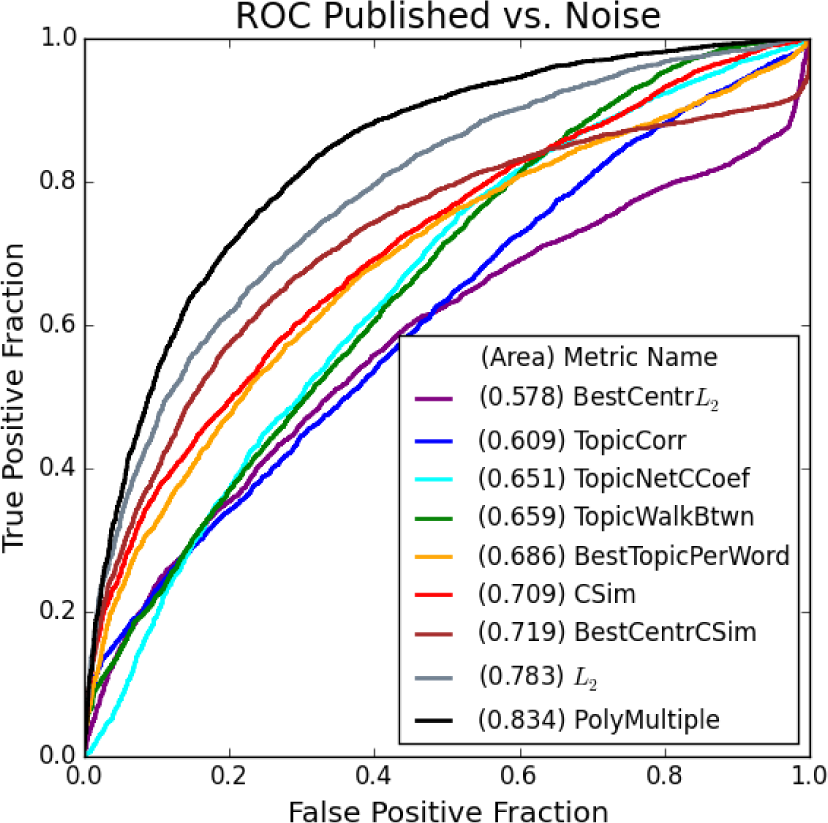
The above ROC curves show the ability for each of our proposed methods to distinguish the MOLIERE results of published pairs from noise. We use our system to generate hypotheses regarding 8,638 pairs, half from each set, on publicly available data released prior to 2,015. We only show the best performing metrics from Section 4.5 for clarity.

**Figure 5:**
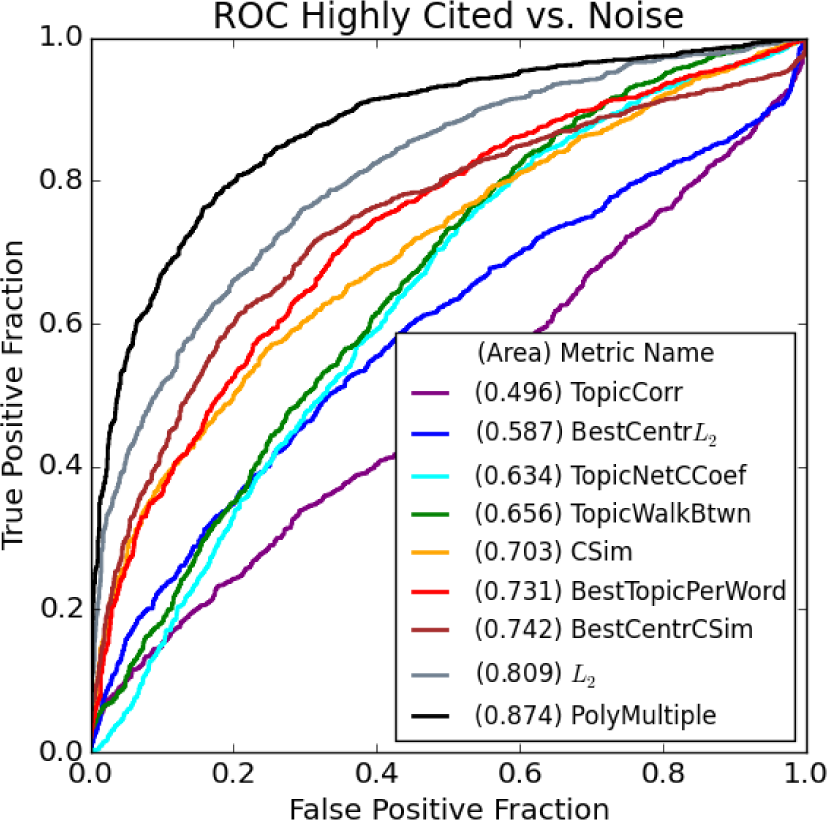
The above ROC curves show the ability for each of our proposed methods to distinguish the MOLIERE results of highly-cited pairs from noise. We identify 1,448 pairs who first occur in papers with over 100 citations published after our cut date. To plot the above ROC curve, we also select an random subset of equal size from the noise pairs.

**Table 2:**
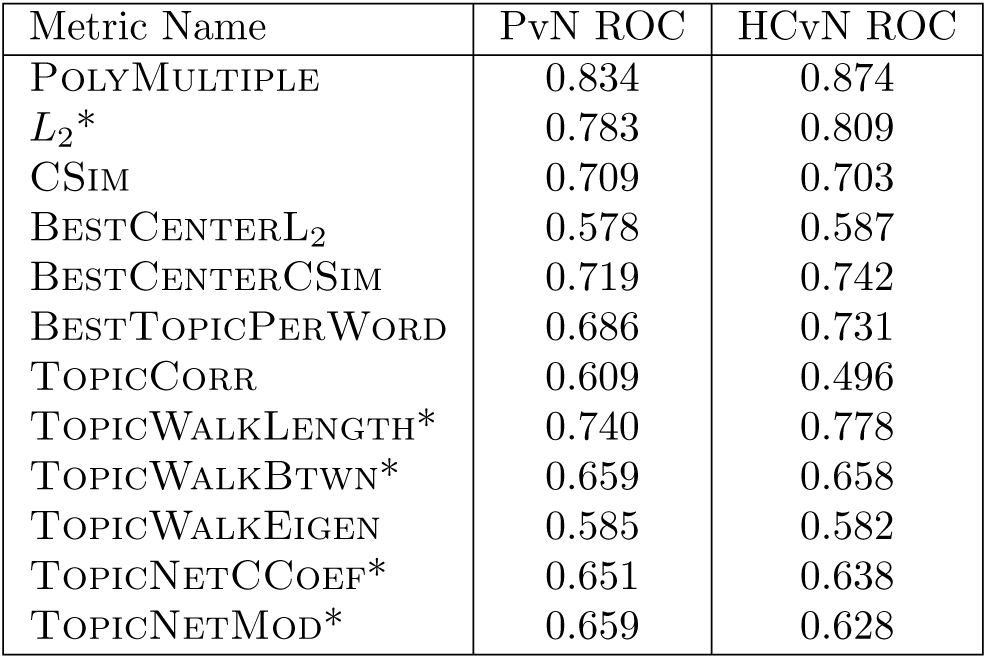
The above summarizes all ROC area results for all considered metrics on the set of published vs. noise pairs (PvN) and highly-cited vs. noise pairs (HCvN). Note, we report areas in the range [0.5,1], so metrics marked with a (*) have been sorted in reverse order for the ROC calculations. This indicates that a higher score along a starred metric indicates a noise pair.

**Topic Model Correlation** metric (see Section 4.2) is a poorly performing metric with an ROC area of 0.609 (PvN) and 0.496 (HCvN). The core issue of this method is its sensitivity to the number of topics generated, and given that we generate 100 topics per pair, we likely drive down performance through topics which are unrelated to the query. Surprisingly, this metric is less able to distinguish highly-cited pairs, which we suppose is because highly-cited connections often bridge very distant concepts [35] and likely results in more noisy topic models. Additionally, we may be able to limit this noise by tuning the number of topics returned from a query, as described in Section 7.

***L*_2_-based metrics** exhibit even more surprising results. BESTCENTRL_2_ performs poorly, with an ROC area of 0.578 (PvN) and 0.587 (HCvN), while the much simpler *L*_2_ metric is exceptional, scoring a 0.783 (PvN) and 0.809 (HCvN). We note that if two words are related, they are more likely to be closer together in our vector space. We evaluate topic centroids based on their closeness to the midpoint between *a* and *c*, normalized by the distance between them, so if that distance is small, the radius from the midpoint is small as well. Therefore, it would seem that the distance between *a* and *c* is a better connection indication, and that the result of the centroid measurement is worse if this distance is small.

**CSim-based metrics** are more straight-forward. The simple CSIMmetric scores a 0.709 (PvN) and 0.703 (HCvN), which is interestingly consistent given that the *L*_2_ metric increases in ROC area given highly-cited pairs. The BESTTOPICPERWORD metric only scores a 0.686 (PvN), but increases substantially to 0.731 (HCvN). The topic centroid method BESTCENTROIDCSIM is the best cosine-based metric with an ROC area of 0.719 (PvN) and 0.742 (HCvN). This result is evidence that our initial hypothesis described in Figure 2 holds given cosine similarity, but as stated above, does not hold for euclidean distance.

**Topic network** metrics are all outperformed by simple *L*_2_, but we see interesting properties from their results that also help users to interpret generated hypotheses. For instance, we see that TOPICWALKBETWEENESS is a negative indicator while TOPICWALKEIGEN is positive. Looking at the example in Figure 3 we see that *a* and *c* are both far from the center of the network, connected to the rest of the topics through a very small number of high-betweenness nodes. In contrast, we see that in the network created from a published pair, the path from *a* to *c* is more central. We also see a denser clustering for the noise pair network, which is echoed by the fact that TOPICNETCCOEF and TOPICNETMODULATIRY are both negative indicators. Lastly, we see that TOPICWALKLENGTH performs the best out of these network approaches, likely because it is most similar to the simple *L*_2_ or CSIM metrics.

**Combination of metrics,** POLYMULTIPLE, significantly outperforms all others with ROC areas of 0.834 (PvN) and 0.874 (HCvN). This is unsurprising because each other metric makes a different assumption about what sort of topic or vector configuration best indicates a published pair. When each is combined, we see not only better performance, but their relative importances. Looking into Table 1 we see that the two *L*_2_-based metrics are most important, followed by the topic network methods, and finally by TOPICWALKCORR and BESTTOPICPERWORD. Unsurprisingly, the coefficient signs correlate directly with whether each metric is a positive or negative indication as summarized in Table 2. Additionally, the ordering of importance roughly follows the same ordering as the ROC areas.

## 6 Related Work and Proposed Validation

The HG community struggles to validate its systems in a number of ways. Yetisgen-Yildiz and Pratt, in their chapter “Evaluation of Literature-Based Discovery Systems,” outline four such methods (M1-M4) [7, 55].

**M1: Replicate Swanson’s Experiments.** Swanson, during his development of ARROWSMITH [39], worked alongside medical researchers to uncover a number of new connections. These connections include the link between Raynaud’s Disease and Fish Oil [45], the link between Alzheimer’s Disease and Estrogen [38] and the link between Migraine and Magnesium [47]. As discussed in [55], a number of projects have centered their validation effort around Swanson’s results [16, 26, 11, 5, 53, 44, 19, 32]. These efforts always rediscover a number of findings using information before Swanson’s discovery date, and occasionally apply additional metrics such as precision and recall in order to quantify their results [14].

While limiting discussion to Swanson’s discoveries reduces the domain of discovery drastically, at its core this method builds confidence in a new system through its ability to find known connections. We expand on this idea by validating automatically and on a massive scale, freeing our discourse from a single researcher’s findings.

**M2: Statistical Evaluation.** Hristovski et al. validate their system by studying a number of relationships and note their confidence and support with respect to the MEDLINE document set [18]. Then, they can generate potential relationships for the set of new connections added to UMLS [1] or OMIM [13]. By limiting their method to association rules, Hristovski et al. note that they can validate their system by predicting UMLS connections using data available prior to their publications. Therefore, this method is the similar to our own, but we notice that restricting discussion to only UMLS gene-disease connections results in a much smaller set than the predicate information present with SemMedDB.

Pratt et al. provide additional statistical validation for their system LitLinker [32]. This method also calculates precision and recall, but this time focusing on their *B*-set of returned results. Their system, like ARROWSMITH [39], returns a set of intermediate terms which may connect two queried entities. Pratt et al. run LitLinker for a number of diseases on which they establish a set of “gold standard” terms. Their method is validated based on its ability to list those gold-standard terms within its resulting *B-sets. This approach requires careful selection of a (typically small) set of gold-standard terms, and is limited to “ABC” systems like ARROWSMITH, which are designed to identify term lists* [36].

**M3: Incorporating Expert Opinion.** This ranges from comparisons between system output and expert output, such as the analysis done on the Manjal system [44], to incorporating expert opinion into gold-standard terms for LitLinker [32], to running actual experiments on potential results by Wren et al. [54]. Expert opinion is at the heart of many recent systems [51, 49, 27, 10, 46, 2], including the previous version of our own. This process is both time consuming and risks introducing significant bias into the validation.

Spangler incorporates expert knowledge in a more sophisticated manner through the use of visualizations [42, 43]. This approach centers around visual networks and ontologies produced automatically, which allows experts to see potential new connections as they relate to previously established information. This view is shared by systems such as DiseaseConnect [27] which generates sub-networks of ONIM and GWAS related to specific queries. Although these visualizations allow users to quickly understand query results, they do not lend themselves to a numeric and massive evaluation of system performance.

BioCreative is a set of challenges focused on assessing biomedical text mining, is the largest endeavor of its kind, to the best of our knowledge [17]. Each challenge centers around a specific task, such as mining chemical-protein interactions, algorithmically identifying medical terms, and constructing causal networks from raw text. Although these challenges are both useful and important, their tasks fall under the umbrella of *information retrieval* because they compare expert analysis with software results given the same text.

**M4: Publishing in the Medical Domain.** This method is exceptionally rare and expensive. The idea is to take prevalent potential findings and pose them to the medical research community for another group to attempt. Swanson and Smalheiser use this technique, which solidifies many of their most prevalent findings, but we are not aware of any other group to attempt this.

An alternative to this idea is taken by Soldatova and Rzhetsky wherein a “robot scientist” automatically runs experiments posed by their HG system [40, 34]. This system uses logical statements to represent their hypotheses, so new ideas can be posed through a series of implications. Going further, their system even identifies statements that would be the most valuable if proven true [35]. *But, the scope of experiments that a robot scientists can undertake is limited; in their initial paper, the robot researcher is limited to small-scale yeast experiments. Additionally, many groups cannot afford the space and expense that an automated lab requires.*

## 7 Deployment Challenges and Open Problems

**Challenge Size.** Our proposed validation challenge involves ranking millions of published and noise query pairs. But, in Section 5 we show our results on a randomly sampled subset of our overall challenge set. This was necessary due to performance limitations of MOLIERE, a system which initially required a substantial amount of time and memory to process even a single hypothesis. To compute these results, we ran 100 instances of MOLIERE, each on a 16 core, 64 GB RAM machine connected to a ZFS storage system. Unfortunately, performance limitations within ZFS created a bottleneck that both limited our results and drastically reduced cluster performance overall. So, our results represent a set of predicates that we evaluated in a limited time period.

**System Optimizations.** While performing a keyword search, most network-centered systems are either I/O or memory bound simply because they must load and traverse large networks. In the case of MOLIERE, we initially spent hours trying to find shortest paths or nearby abstracts. But, we found a way to leverage our embedding space and our parallel file system in order to drastically improve query performance.

In brief, one can discover a relevant knowledge-network region by inducing a subnetwork on the set of keywords found within the hyperellipse focused between *a* and *c*. This increases performance because, given a parallel file system, creating an edge list for that subnetwork is a massively parallel task of order *O*(*n/p*). Additionally, when finding a path from *a* to *c* in that subnetwork, we can leverage the embeddings again to replace our shortest-path algorithm with *A*\*. We note that if such a path does not exist within our subnetwork, one can always increase the hyperellipse constant in order to broaden the subnetwork.

The overall effect of our optimizations reduced the wall-clock runtime of a single query from about 12 hours to about 5-7 minutes. Additionally, we reduced the memory requirement for a single query from over 400GB to under 16GB. Moreover, because the quality of our system’ss results is tunable through the hyperellipse constant, we note that our results presented here are nearly identical to our previous software.

**Highly Cited Predicates.** Identifying highly sighted predicates requires that we synthesize information across multiple data sources. Although SemMedDB contains MEDLINE references for each predicate, neither contains citation information. For this, we turn to Semantic Scholar because not only do they track citations of medical papers, but they allow a free bulk download of metadata information (many other potential sources either provide a very limited API or none at all). In order to match Semantic Scholar data to MEDLINE citation, it is enough to match titles. This process allows us to get citation information for many MEDLINE documents, which in turn allow us to select predicates who’s first occurrence was in highly cited papers.

We explored a number of thresholds for what constitutes “highly cited” and selected 100 because it was a round number that provided a sizable number of selected predicates. Because paper citations approximately follow a power-law distribution, any change would have drastically changed the size of this set. We note that the set of selected predicates was also limited by the quality of data in Semantic Scholar, and that the number of citations identified this was appeared to be substantially lower than that reported by other methods. Nevertheless, because our available citation count is a lower bound of the real value, we are assured that our “highly cited” set is actually well cited. According to our observations the least cited paper was cited more than 400 times in Google Scholar.

**Quality of Predicates.** Through our above methods we learned that careful ranking methods can distinguish between published and noise predicates, but there is a potential inadequacy in this method. Likely, there exist a number of predicates within our published set which despite occurring in an abstract, are untrue. Additionally, it is possible that a noise predicate may be discovered to be true in the future. If our system ranks the published predicate which is untrue below the noise predicate which is, we worsen our system’s ROC area. This same phenomena is addressed by Yetisgen-Yildiz and Pratt when they discuss the challenges present in validating literature-based discovery systems [55] — if a HG systems goal is to identify novel findings, then it *should* find different connections than human researchers.

We show through our results that despite an uncertain validation set, there are clearly core differences between publishable results and noise, which are evident at scale. Although there may be some false positives or negatives, we see through our meaningful ROC curves that they are far outnumbered by more standard predicates.

**Automatic Question Posting.** Going forward we wish to study highly ranked noise predicates for the purpose of automatic question posing. This would mean that our system would search through its set of entities, run queries, and report the most promising potential new connections. In order to do this effectively we need to gain an understanding of how we can intelligently search local regions of our knowledge network and how to define locality for this task.

**Comparison with ABC Systems.** Additionally, we would like to explore how our ranking methods apply to traditional ABC systems. Although there are clear limitations to these systems [36], many of the original systems such as ARROWSMITH follow the pattern. These systems typically output a list of target terms and linking terms, which could be thought of as a topic. If we were to take a pretrained embedding space, and treated a set of target terms like a topic, we could likely use our methods from Section 4 to validate any ABC system.

**Verb Prediction.** We noticed, while processing SemMedDB predicates, that we can improve MOLIERE if we utilize verbs. SemMedDB provides a handful of verb types, such as “TREATS,” “CAUSES,” or “INTERACTS_WITH,” that suggest a concrete relationship between the subject and object of a sentence. MOLIERE currently outputs a topic model that can be interpreted using our new metrics, but does not directly state what sort of connection may exist between *a* and *c*. So, we want to see if we can accurately predict these verb types given only topic model information.

**Interpretability of hypotheses** remains one of the major problems in HG systems. Although topic-driven HG partially resolve this issue by producing a readable output, we still observe many topic models *T* (i.e., hypotheses) whose *T_i_* ∈ *T* are not intuitively connected with each other. While the proposed ranking is definitely helpful for understanding *T*, it still does not fully resolve the interpretability problem. One of our current research directions to tackle it is using text summarization techniques.

## 8 Conclusion

When we rely on experts to pose questions or perform experiments for the sake of validation, we drastically reduce the productivity of all involved. This is compounded by the need for software systems to rapidly iterate over time, gaining features and trying new methods to produce results. That work flow cannot afford the time of experts. Although we are not the first to identify the need for large-scale validation, we do propose a validation method which is impartial, does not need expert opinion, and operates at large scale. Furthermore, by selecting different cut years, we can even continue with our validation scheme long into the future, without manually selecting discoveries.

Additionally, our validation challenge can be taken by any system, not just an ABC system, so long as that system can rank its results. As we show in Section 4, even topic-driven systems can be adapted to produce such a ranking. Our validation methodology even extends across domains, as long as one can extract predicates. Overall, we see an immense need for validation which allows us to meaningfully compare performance across HG systems, or across scientific domains, and we are unaware of any other methodology for doing so.

To aid the biomedical HG community, we provide our code, query sets, and training data on-line at bit.ly/2EtVshN. With these resources, anyone could replicate our results using MOLIERE, or incorporate our ranking methods or validation framework into their own HG system.

